# Williams Syndrome-Specific Neuroarchitectural Profile and Its Associations with Cognitive Features

**DOI:** 10.1101/060764

**Authors:** Chun Chieh Fan, Timothy T. Brown, Hauke Bartsch, Joshua M. Kuperman, Donald J. Hagler, Andrew Schork, Eric Halgren, Anders M. Dale

## Abstract

Williams Syndrome (WS), a rare genetic disorders caused by hemizyous deletion of ~26 genes on the chromosome 7, has unique cognitive features and neuroanatomic abnormalities. Limited in statistical power due to its rareness had led to inconsistent in many direct comparisons using structural magnetic resonance imaging (MRI), and their associations with cognitive features of WS are not clear. Here, we used a novel approach to derive a WS specific neuroarchitectural profile and tested its association with cognitive features of WS. Using a WS adult cohort (n = 43), we trained a logistic elastic-net model to extract a sparse representation of WS specific neuroarchitectural profile. The predictive performances are robust within the training cohort (leave one out cross-validation AUC = 1.0) and generalized well in an independent teenager WS cohort (n = 60, AUC = 1.0). The WS specific neuroarchitectural profile includes multiple MRI measurements in the orbitofrontal cortex, superior parietal cortex, Sylvian fissures, and basal ganglia, whereas its variations reflect the underlying size of hemizygous deletion, and mediated the disease impact on the cognitive features of WS. In this study, we demonstrate the robustness of the derived WS specific neuroarchitectural profile, suggesting the joint developmental abnormalities in the cortical-subcortical circuitry cause the unique features of WS cognition.

## Introduction

Williams Syndrome (WS) is a rare multi-systemic disorder caused by deletion of ~26 genes on the chromosome 7. Its unique cognitive features, including intellectual impairment, visuospatial deficits, and hypersociability (Pober, 2010), has led to extensive researches using magnetic resonance imaging (MRI), in a hope to identify the intermediating neural processes from genetic deletions to cognitive impact (Martens *et al.*, 2008). Previous MRI studies had found, what distinguish WS from other genetic disorder with intellectual impairment, e.g. Down syndrome, is not the reduced total brain volumes per se, but the aberrant regionalization of brain (Jernigan and Bellugi, 1990). However, it is still poorly understood which regionalization patterns are specific to WS and related to its underlying genetic deletions. The most consistent findings are the gyral patterns in the superior parietal regions and orbital frontal cortex, which were repeatedly found to be different between WS patients and healthy individuals (Meyer-Lindenberg *et al.*, 2004, Kippenhan *et al.*, 2005, Gaser *et al.*, 2006). Yet their specificity to WS and relevance to the WS distinct cognitive features were left unanswered. Abnormalities in the Sylvian fissures (Eckert *et al.*, 2006) and reduced volumes of subcortical structures (Meyer-Lindenberg *et al.*, 2006) were also reported, yet not consistently found in other WS cohorts (Meda *et al.*, 2012). Moreover, none had shown whether the variations of these abnormalities reflect the size of genetic deletion in the WS related chromosomal regions. The lack of evidence might be due to limited statistical power in previous studies. The rarity of patients with WS, and even more rare prevalence for patients with partial deletions in the WS related chromosomal regions makes the quantitative comparisons across MRI measures and groups impractical.

Here, we re-examined the WS specific neuroarchitectural profile with new analyzing strategy. We utilized the characteristics of different cohorts in order to maximize the statistical power. First, we extracted the WS specific neuroarchitectural profile from an adult WS cohort. To deal with the large amount of MRI measures and limited sample size, we fit an elastic-net logistic regression to find the best predictive and sparse features. The resulting model then provides the basis for calculating WS neuroanatomic scores that represent the similarity of individuals’ brain to WS given their MRI measures. The generalizability of the WS specific neuroarchitectural profile was tested in an independent teenager WS cohort. After establishing the generalizability of the model, we examined whether the variations of the scores could reflect the reduced size of genetic deletions in WS chromosomal regions and were associated with cognitive features of WS.

## Materials and Methods

Two independent cohorts were included in the current analysis (Table 1). The first cohort consisted of 42 adult participants in total, including adult patients with WS, patients with atypical deletions in WS related chromosomal regions, and healthy controls (HC). We used the adult cohort to extract the WS-specific neuroarchitectural profile. The generalizability of the WS-specific neuroarchitectural profile was tested in an independent cohort, consisting of 23 typically developing teenagers (TD) and 37 individuals with heterogeneous diagnoses, including WS, high functioning autism (HFA), specific language impairment (SLI), and focal lesions in the brain (FL). The demographic characteristics of each cohort are shown in Table 1. The following section describes the recruitment, measurements, and MRI protocols for each cohort.

**Table 1.** Demographics and global MRI measurements of participants in two cohorts.

| Groups | n | Age – years | Gender – Male | Full IQ | SISQ – AS | SISQ – ES |
| --- | --- | --- | --- | --- | --- | --- |
| **Adult WS cohort** |  |  |  |  |  |  |
| WS | 22 | 31.6 (10.8) | 59% | 66.6 (5.0) | 5.4 (1.4) | 5.8 (0.8) |
| HC | 16 | 25.9 (7.0) | 37% | 96.7 (14.6) | 3.6 (1.2) | 4.4 (1.0) |
| Atypical WS | 5 | 17.7 (2.5) | 20% |  |  |  |
| **Teenager cohort** |  |  |  |  |  |  |
| WS | 7 | 11.95 (1.75) | 29% |  |  |  |
| TD | 23 | 9.48 (1.87) | 52% |  |  |  |
| FL | 8 | 9.73 (1.26) | 50% |  |  |  |
| HFA | 14 | 9.83 (1.45) | 79% |  |  |  |
| SLI | 8 | 10.09 (1.48) | 75% |  |  |  |

WS: Williams Syndrome. HC: Healthy controls. TD: Typical developed individuals. FL: Individuals with focal lesions in the MRI scans of brain. HFA: high function autism. SLI: specific language impairment.

### *Adult WS cohort*

The adult cohort consisted of 22 typical WS patients, 5 atypical WS patients, and 16 HC. Part of this cohort has been involved in a series of MRI studies for WS that were published elsewhere (Eckert *et al.*, 2006, Van Essen *et al.*, 2006). In short, classification of WS was based on clinical presentation and genetic criteria using fluorescent in-situ hybridization. The typical WS patients were defined as those who showed a deletion of all ~26 genes in the 7q11.23 region. Patients with atypical WS have a shorter hemizygous deletion on the WS-related chromosomal regions. HCs were screened for a history of neurological disorders, psychiatric illness, and substance uses. Participants’ intelligence was assessed with the Wechsler Intelligence Scale 3^rd^ Edition (Wechsler, 2008). Sociability was assessed with the Salk Institute Sociability Questionnaire (SISQ) (Doyle *et al.*, 2004). All participants were scanned with a 1.5 Tesla MRI scanner (GE Signa TwinSpeed scanner, echo time (TE) = 3.0 msec, repetition time (TR) = 8.7msec, inversion time (TI) = 270 msec, flip angle (FA) = 8°, field of view (FoV) = 24 cm, voxel size = 1.25 × 1.25 × 1.2 mm). To correct for possible motion artifacts, real-time prospective motion tracking and correction (PROMO) was used for all participating subjects (White *et al.*, 2010). Distortions caused by nonlinearity of the spatial encoding gradient fields were corrected with predefined nonlinear transformations (Jovicich *et al.*, 2006). Non-uniformity of signal intensity was reduced with the nonparametric nonuniform intensity normalization method (Sled *et al.*, 1998).

### Teenager cohort

The teenager cohort in the current analysis consisted of 7 WS patients and 53 non-WS individuals, ranging from 6 to 13 years of age (Table 1). Detailed recruiting procedures and diagnostic criteria can be found in previously published studies (Mills *et al.*, 2013). In brief, teenagers with WS were diagnosed using the same clinical presentation and genetic criteria mentioned in the previous section. Subjects in the TD group were recruited from the community, meeting criteria including normal performance on the standardized language test, normal intelligence, and no history of developmental or language delay. Individuals with HFA, SLI, and FL were diagnosed and recruited from populations at a local pediatric neurology clinic and a clinic for speech and language disorders (Mills *et al.*, 2013). Considering the purpose of this analysis was to determine whether the extracted brain features can be used to distinguish children with WS from a heterogeneous group, we pooled the TD, HFA, SLI, and FL subjects into one group as non-WS in the following analysis. All participants were scanned with MRI using the identical imaging protocol that was used with the adult WS cohort.

### Imaging Data Processing

After initial image data inspection and quality control, T1-weighted images underwent automated processing using methods implemented in Freesurfer software (Dale *et al.*, 1999, Fischl *et al.*, 1999). This automated processing corrects variations in image intensity due to RF coil sensitivity inhomogeneities, registered to a common reference, and then segments volumes into cortical and subcortical structures. Four different morphological measures of T1-weighted images were derived, including the volumes of subcortical structures (Dale *et al.*, 1999), sulcal depths of the cortical surface (Fischl *et al.*, 1999), cortical surface area (Chen *et al.*, 2012), and geometric deformations of the cortical surface (Fan *et al.*, 2015). The first measure comes from the segmentation step while the remaining measures require further cortical surface reconstruction involving surface tessellation and spherical mapping to ensure comparability across subjects (Fischl *et al.*, 1999). Sulcal depth is the distance from each point on the cortical surface to the average mid plane of the cortical surface, capturing the gyrification of the brain. Cortical surface area expansion is the neighboring area of a given cortical surface point divided by the total surface area. The geometric deformation is the 3D Cartesian coordinates of the cortical surface, characterizing the folding patterns of the brain. Subcortical structure volumes were divided by total brain volumes, and sulcal depths and geometric deformations were divided by the cubic root of each total brain volume to produce a uniform index as well as to control for the global brain volume differences.

### Statistical Analysis

To characterize WS specific neuroarchiectural profile from multiple MRI measures and determine their relative importance, we fit an elastic-net logistic regression using data from adult cohort and check their performance with leave-one-out cross validation (LOOCV). The index for model performance was area under curve (AUC) in the ROC analysis. The model included all four MRI measures, using the ridge penalties to reduce the problem of unknown correlations and additional lasso penalties for selecting best predictive features. The tuning parameters were optimized during the cross-validation. After deriving the WS specific neuroarchitectural profile from previous training step, the model was applied to the teenager cohort to see if the model can predict WS status out of a heterogeneous pool. We also applied to the adult patients with atypical deletion size in the WS related chromosomal regions to check if the model can reflect the underlying reduced deletion size in the WS related chromosomal regions. Afterwards, the relationships between model-predicted scores and cognitive measures were explored using mediation analysis. Sobel tests were used to examine whether the group differences were mediated by the neuroarchitectural profile. Pearson correlations were further conducted in adult cohort within each group to see if the predicted scores associated with within-group variations of cognitive function.

## Results

### Deriving the profile and its generalizability

In the discriminant analysis with LOOCV, the AUC of the WS specific neuroarchitectural profile achieved 1.00 in the adult WS cohort (two tail test for AUC greater than 0.5, p < 0.05). The model removes 98.4 percent of the input variables, leaving 412 variables from four MRI measures. Among individuals with reduced size of deletions in the WS related chromosomal regions, their predicted scores of the WS specific neuroarchitectural profile are significantly higher than HC (t_19_ = 9.4, p < 10^-7^), while lower than patients with typical WS (t_25_=-2.2, p = 0.038). To further test the generalizability of the model, we applied the WS specific neuroarchitectural profile to a teenager cohort with heterogeneous developmental status. In this independent cohort, the WS specific neuroarchitectural profile retains high specificity to WS, achieving AUC with 1.0 (Figure 1).

**Figure 1.**
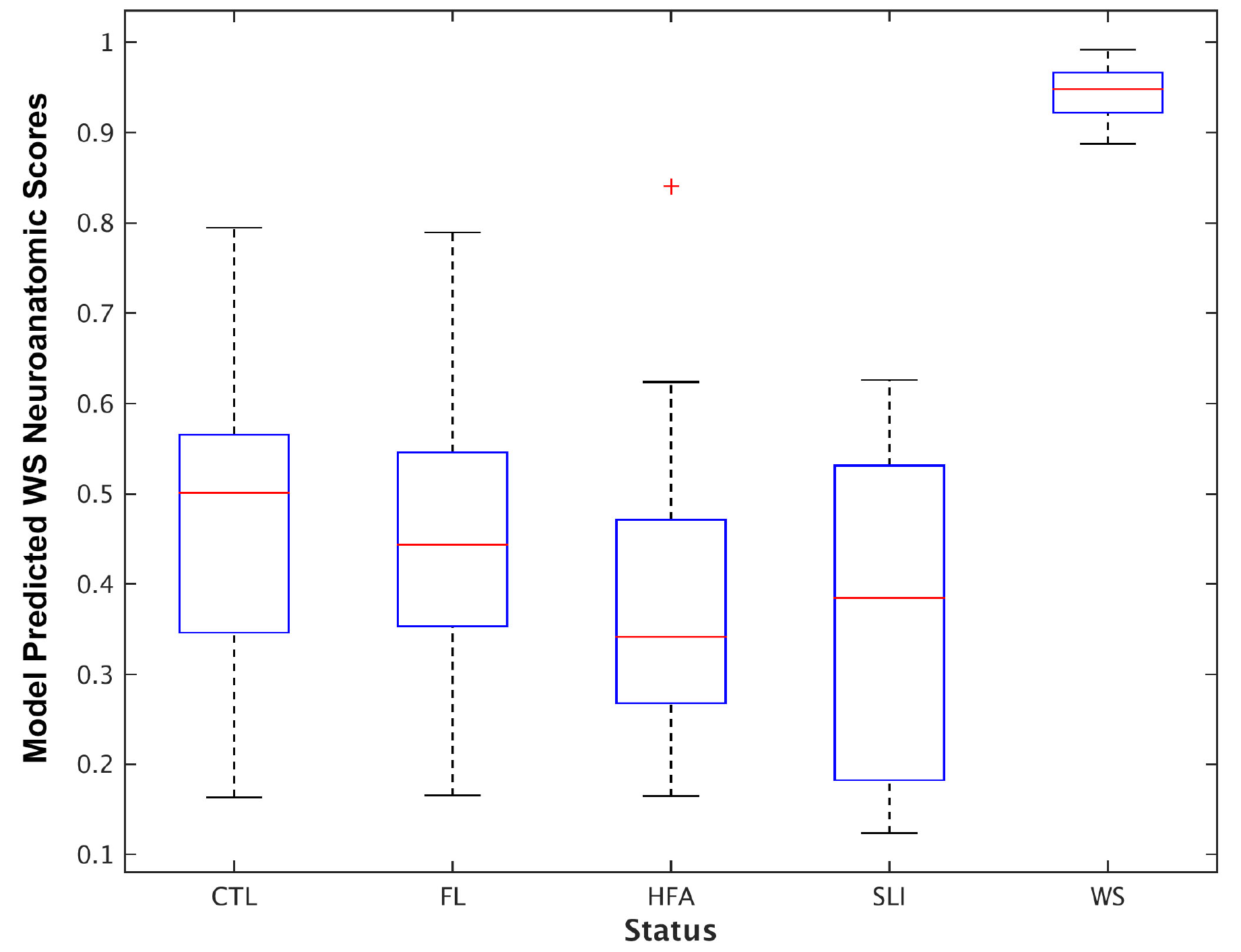
Boxplot of model predicted WS neuroanatomic scores across groups in the teenager cohort.

### Features in the WS specific neuroarchitectural profile

The cortical surface features extracted by the elastic-net model were shown in the Figure 2. Since the input variables are normalized, the weights of selected features reflect the relative importance for predicting WS. Selected local features can be observed across cortical surface regions, yet sparing the dorsal and medial part of frontal cortex. Orbitofrontal cortex and superior parietal cortex contain predictive features consistently across all three cortical surface measures (Figure 2). In addition, cortical surface area contained predictive features in the Sylvian fissure and temporal poles. Two subcortical structures were also selected. Disproportionally decreasing sizes of left putamen (weights = −0.010) and left nucleus accumbens (weights = −0.014) were predictive for WS status.

**Figure 2.**
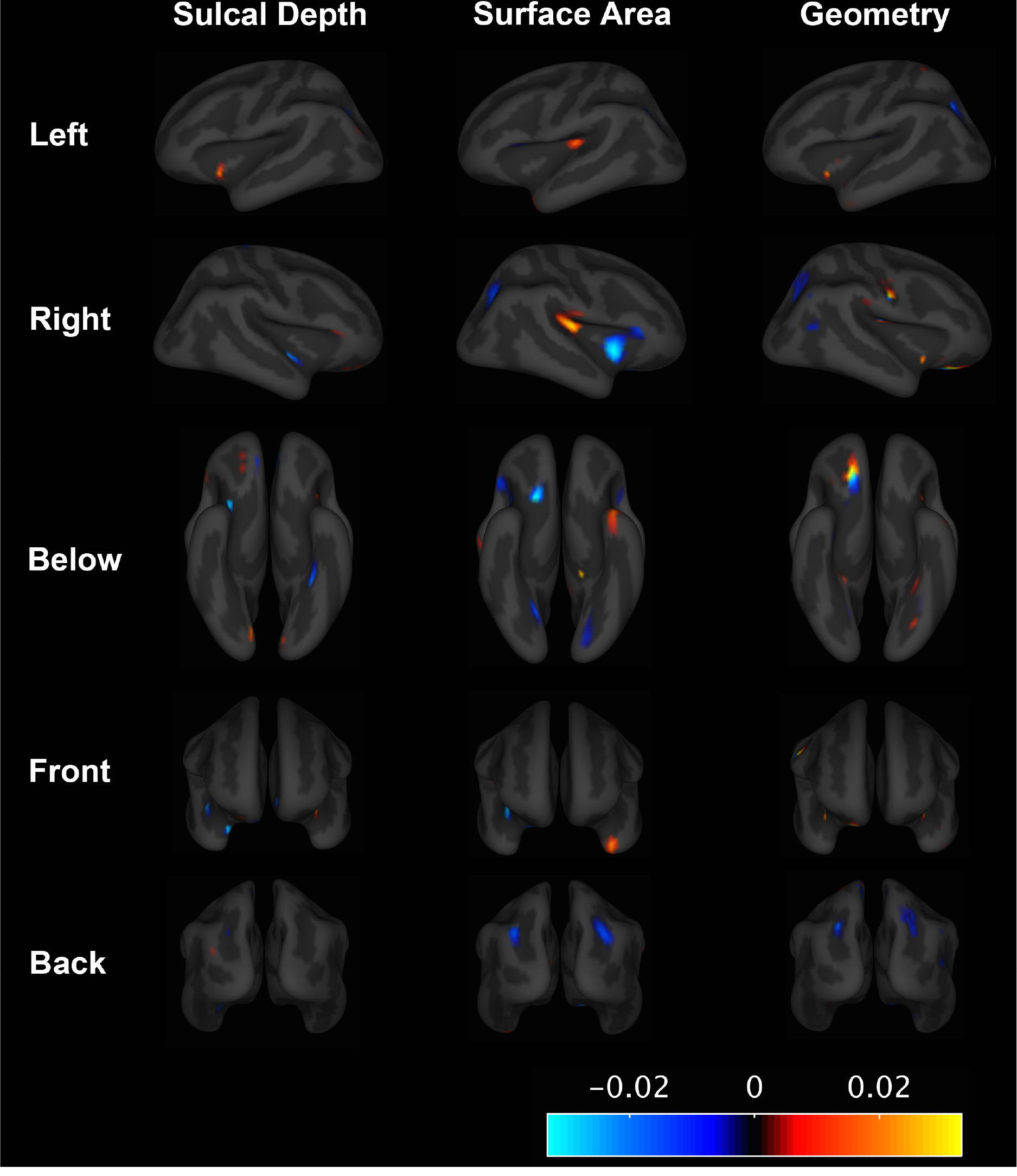
Elastic net model learnt features for predicting WS status.

### Associations with WS cognitive features

The relationships among WS status, the WS specific neuroarchitectural profile, and cognitive function of WS were illustrated in the Table 2. The Sobel tests for mediation indicate that the group differences in general intelligence, SISQ stranger score, and SISQ empathy score are largely explained by the mediating effect of the WS specific neuroarchitectural profile (all p values < 10^−3^, Bonferroni correction). In the within-group analyses, the variations of the WS specific neuroarchitectural profile are still significantly associated with SISQ empathy score, though in trend p-values after Bonferroni correction for 9 independent tests (p = 0.063).

## Discussion

Our study is the first to use a multidimensional imaging approach to characterize the WS-specific neural architectural features. Orbitofrontal cortex, superior parietal cortex, and Sylvian fissures differed significantly across imaging measures (Figure 2). Brain volumes were disproportionally reduced in the putamen and nucleus accumbens. Summarizing those observed differences, our extracted WS-specific neural architectural profile of features robustly predicted the WS status in both adult and teenager cohorts (Figure 1). It also demonstrated a robust dosage effect of the hemizygous deletions within the WS-related chromosomal regions, and was associated with one of the cardinal cognitive features of WS (Table 2).

**Table 2.** Mediating effects and within group correlations between model predicted WS neuroanatomic scores and cognitive measures.

|  | Mediating Effect* |  | Within HC |  | Within WS |  |
| --- | --- | --- | --- | --- | --- | --- |
| FIQ | z = -6.31 | p = 1e-10 | r = 0.29 | p = 0.34 | r = 0.18 | p = 0.52 |
| SISQ – Stranger | z = 3.73 | p = 9e-5 | r = -0.01 | p = 0.96 | r = 0.10 | p = 0.74 |
| SISQ - Empathy | z = 4.61 | p = 2e-6 | r = 0.09 | p = 0.74 | r = 0.70 | p = 7e-3 |

\* The mediating effect is checked with Sobel test for mediation, treating model predicted WS neuroanatomic scores as the mediator and each cognitive measure as dependent variable.

The observed differences in cortical surface measures are consistent with previous reports (Kippenhan *et al.*, 2005, Eckert *et al.*, 2006, Gaser *et al.*, 2006, Meyer-Lindenberg *et al.*, 2006, Van Essen *et al.*, 2006, Martens *et al.*, 2008). Gyrification abnormalities in the orbitofrontal cortex, Sylvian fissures, and superior parietal regions have been repeatedly reported (Kippenhan *et al.*, 2005, Eckert *et al.*, 2006, Gaser *et al.*, 2006, Van Essen *et al.*, 2006). Some have hypothesized that the gyrification differences in WS patients result from reduced arealization of the cortical surface (Gaser *et al.*, 2006, Van Essen *et al.*, 2006). Our findings are the first to demonstrate that WS patients indeed show reduced cortical surface area in the orbitofrontal and superior parietal regions (Figure 2). The Sylvian fissure abnormalities might result from a complex interaction between the area expansion of temporal parietal junction and reduced arealization of insular regions (Figure 2). In general, these observed differences match with the cognitive profile of WS patients. The superior parietal regions has been linked to the visuospatial defect of WS (Meyer-Lindenberg *et al.*, 2004) while the temporal parietal junction, insula, and orbitofrontal cortex have been associated with socially relevant functions (Adolphs, 2001, Saxe and Kanwisher, 2003, Meyer-Lindenberg *et al.*, 2005).

In terms of subcortical volumes, previous reports are conflicting. Some have found that amygdala volumes are disproportionally reduced but other studies have not (Meyer-Lindenberg *et al.*, 2005, Haas *et al.*, 2010). Our results suggest that amygdala volumes remain relatively in proportion to total brain volume. This finding supports the notion that cortical abnormalities rather than the deficits in the amygdala lead to aberrant cortical-amygdala functional pairing in WS (Meyer-Lindenberg *et al.*, 2005). On the other hand, disproportionate differences in the putamen suggest that the WS pathology might involve other frontal subcortical circuitry. Yet, currently there is no clear evidence to suggest which of the frontal-subcortical circuitry is mechanistically responsible. Studies using diffusion imaging might be helpful to further clarify this issue (Marenco *et al.*, 2007).

The neural architectural differences between WS patients and controls are not limited to only those regions highlighted above. Small sample sizes are common in published studies of WS, considering the prevalence of WS is very rare (Pober, 2010). Our approach for extracting WS-specific features circumvents this limitation of group comparisons. The extracted WS-specific profile of features robustly identified patients with WS in all scenarios while also being significantly associated with cognitive features of WS (Table 2).

Previous case studies have indicated that atypical WS patients with smaller genetic deletions have lower sociability than typical WS patients (Doyle *et al.*, 2004). Duplications of the genes in the WS-related chromosomal regions increase the risk of autism (Glessner *et al.*, 2009, Mulle *et al.*, 2014). The telomere side of WS-related chromosomal regions, which tends to be spared in smaller deletions, contains genes such as *GTF2I* and *GTF2IRD1,* which have been found to be associated with social behaviors in mouse models (Tassabehji *et al.*, 2005, Young *et al.*, 2008). Our data show a dosage effect of predicted risk scores and that the predicted risk scores are positively correlated with hypersociability. These findings suggest that our extracted WS-specific profile of features might relate directly to the underlying genetic cause of hypersociability.

Taken together, our novel multidimensional imaging approach captures the widespread differences observed within the neural architecture of individuals with WS. It circumvents the limitation of statistical power that are common in many previous studies using only group comparison methods. A major benefit of our analytic strategy is that the extracted features can be readily applied to other imaging datasets. Applications of the extracted features on a large imaging genomic cohort would be helpful for investigating the genetic causes of hypersociability in WS and for exploring the genetic causes of human social behavior in general.

## References

Adolphs R. The neurobiology of social cognition. Current opinion in neurobiology. 2001;11(2):231–9.

Chen CH, Gutierrez ED, Thompson W, Panizzon MS, Jernigan TL, Eyler LT, et al. Hierarchical genetic organization of human cortical surface area. Science. 2012;335(6076): 1634–6.

Dale AM, Fischl B, Sereno MI. Cortical surface-based analysis. I. Segmentation and surface reconstruction. Neuroimage. 1999;9(2):179–94.

Doyle TF, Bellugi U, Korenberg JR, Graham J. “Everybody in the world is my friend” hypersociability in young children with Williams syndrome. American journal of medical genetics Part A. 2004;124A(3):263–73.

Eckert MA, Galaburda AM, Karchemskiy A, Liang A, Thompson P, Dutton RA, et al. Anomalous sylvian fissure morphology in Williams syndrome. NeuroImage. 2006;33(1):39–45.

Fan CC, Bartsch H, Schork AJ, Chen CH, Wang Y, Lo MT, et al. Modeling the 3D geometry of the cortical surface with genetic ancestry. Current biology: CB.2015;25(15):1988–92.

Fischl B, Sereno MI, Dale AM. Cortical surface-based analysis. II: Inflation, flattening, and a surface-based coordinate system. Neuroimage. 1999;9(2):195–207.

Gaser C, Luders E, Thompson PM, Lee AD, Dutton RA, Geaga JA, et al. Increased local gyrification mapped in Williams syndrome. Neuroimage. 2006;33(1):46–54.

Glessner JT, Wang K, Cai G, Korvatska O, Kim CE, Wood S, et al. Autism genome-wide copy number variation reveals ubiquitin and neuronal genes. Nature. 2009;459(7246):569–73.

Haas BW, Hoeft F, Searcy YM, Mills D, Bellugi U, Reiss A. Individual differences in social behavior predict amygdala response to fearful facial expressions in Williams syndrome. Neuropsychologia. 2010;48(5):1283–8.

Jernigan TL, Bellugi U. Anomalous brain morphology on magnetic resonance images in Williams syndrome and Down syndrome. Archives of neurology. 1990;47(5):529–33.

Jovicich J, Czanner S, Greve D, Haley E, van der Kouwe A, Gollub R, et al. Reliability in multi-site structural MRI studies: effects of gradient non-linearity correction on phantom and human data. Neuroimage. 2006;30(2):436–43.

Kippenhan JS, Olsen RK, Mervis CB, Morris CA, Kohn P, Meyer-Lindenberg A, et al. Genetic contributions to human gyrification: sulcal morphometry in Williams syndrome. The Journal of neuroscience: the official journal of the Society for Neuroscience. 2005;25(34):7840–6.

Marenco S, Siuta MA, Kippenhan JS, Grodofsky S, Chang WL, Kohn P, et al. Genetic contributions to white matter architecture revealed by diffusion tensor imaging in Williams syndrome. Proceedings of the National Academy of Sciences of the United States of America. 2007;104(38):15117–22.

Martens MA, Wilson SJ, Reutens DC. Research Review: Williams syndrome: a critical review of the cognitive, behavioral, and neuroanatomical phenotype. J Child Psychol Psyc. 2008;49(6):576–608.

Meda SA, Pryweller JR, Thornton-Wells TA. Regional brain differences in cortical thickness, surface area and subcortical volume in individuals with Williams syndrome. PloS one. 2012;7(2):e31913.

Meyer-Lindenberg A, Hariri AR, Munoz KE, Mervis CB, Mattay VS, Morris CA, et al. Neural correlates of genetically abnormal social cognition in Williams syndrome. Nat Neurosci. 2005;8(8):991–3.

Meyer-Lindenberg A, Kohn P, Mervis CB, Kippenhan JS, Olsen RK, Morris CA, et al. Neural basis of genetically determined visuospatial construction deficit in Williams syndrome. Neuron. 2004;43(5):623–31.

Meyer-Lindenberg A, Mervis CB, Faith Berman K. Neural mechanisms in Williams syndrome: a unique window to genetic influences on cognition and behaviour. Nat Rev Neurosci. 2006;7(5):380–93.

Mills BD, Lai J, Brown TT, Erhart M, Halgren E, Reilly J, et al. Gray Matter Structure and Morphosyntax Within a Spoken Narrative in Typically Developing Children and Children With High Functioning Autism. Developmental neuropsychology. 2013;38(7):461–80.

Mulle JG, Pulver AE, McGrath JA, Wolyniec PS, Dodd AF, Cutler DJ, et al. Reciprocal Duplication of the Williams-Beuren Syndrome Deletion on Chromosome 7q11.23 Is Associated with Schizophrenia. Biological Psychiatry. 2014;75(5):371–7.

Pober BR. Williams-Beuren syndrome. The New England journal of medicine. 2010;362(3):239–52.

Saxe R, Kanwisher N. People thinking about thinking people: the role of the temporo-parietal junction in “theory of mind”. Neuroimage. 2003;19(4):1835–42.

Sled JG, Zijdenbos AP, Evans AC. A nonparametric method for automatic correction of intensity nonuniformity in MRI data. IEEE transactions on medical imaging. 1998;17(1):87–97.

Tassabehji M, Hammond P, Karmiloff-Smith A, Thompson P, Thorgeirsson SS, Durkin ME, et al. GTF2IRD1 in craniofacial development of humans and mice. Science. 2005;310(5751):1184–7.

Van Essen DC, Dierker D, Snyder AZ, Raichle ME, Reiss AL, Korenberg J. Symmetry of cortical folding abnormalities in Williams syndrome revealed by surface-based analyses. The Journal of neuroscience: the official journal of the Society for Neuroscience. 2006;26(20):5470–83.

Wechsler D. Wechsler adult intelligence scale-Fourth Edition (WAIS-IV). San Antonio, TX: NCS Pearson. 2008.

White N, Roddey C, Shankaranarayanan A, Han E, Rettmann D, Santos J, et al. PROMO: Real-time prospective motion correction in MRI using image-based tracking. Magnetic Resonance in Medicine. 2010;63(1):91–105.

Young E, Lipina T, Tam E, Mandel A, Clapcote S, Bechard A, et al. Reduced fear and aggression and altered serotonin metabolism in Gtf2ird1 -targeted mice. Genes, Brain and Behavior. 2008;7(2):224–34.

